# A Charge-reversal Point Mutation Completely Depletes Flavin Chromophore from European Robin Cryptochrome 4a Protein

**DOI:** 10.1101/2025.11.24.690116

**Authors:** Jingjing Xu, Emil Sjulstok Rasmussen, Francis Berthias, Jessica Schmidt, Henrik Mouritsen, Ole N. Jensen, Ilia A. Solov’yov

## Abstract

Cryptochrome 4a (Cry4a) is a magnetically sensitive protein that could enable night-migratory birds to sense the geomagnetic field for navigation. The key to the protein magnetic sensitivity is the flavin adenine dinucleotide (FAD) cofactor, which initiates the electron transfer within the protein leading to a spin-correlated radical pair. Despite its importance, the mechanism of FAD binding in avian Cry4a proteins remains unclear. Here we show that point mutagenesis of positively charged arginine residue at position 356 to negatively charged glutamic acid completely depletes FAD binding from European robin (*Erithacus rubecula*) Cry4a. The result indicates that electrostatic interactions constitute the primary driving force that enables FAD binding in European robin Cry4a. The finding provides new structural insight into the molecular basis of FAD binding in Cry4 and advances our understanding on the biophysical underpinnings of bird magnetoreception.

**Table of Contents Graphic:** 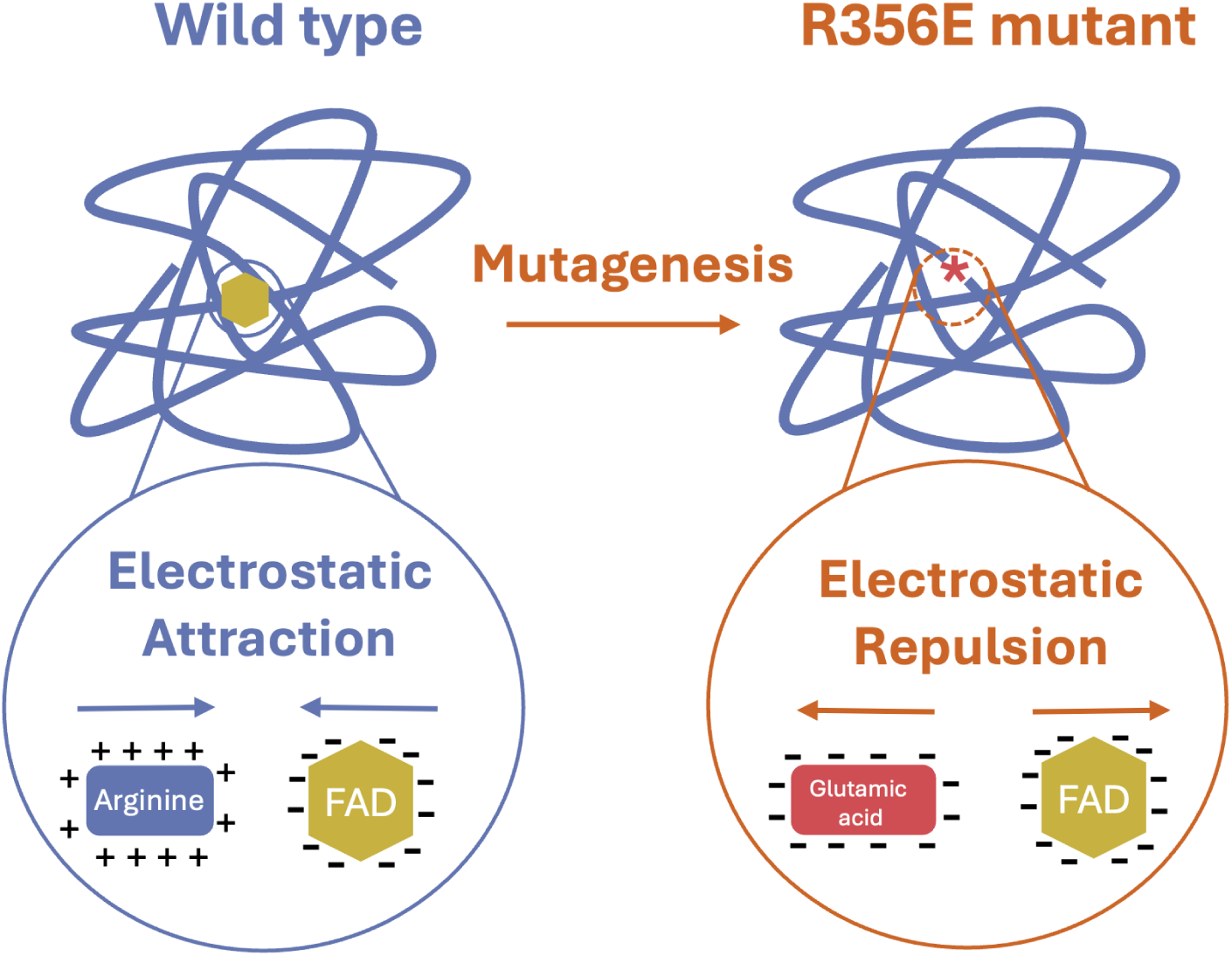

---

Cryptochrome (Cry) proteins are flavoproteins that are involved in diverse biological processes including photoreception, circadian regulation, and potentially magnetoreception. ^1–7^ The diverse biological functions are often associated with the falvin adedine dinucleotide (FAD) cofactor that can be non-covelently bound by some Cry proteins. Essentially, FAD is a blue light absorbing chromophore that can trigger the electron transfer process, leading to a likely biological signaling cascade related to the protein.^8–11^ Animal cryptochromes are classified into Type I, II and IV. Type I Cry proteins, such as *Drosophila melanogaster* Cry (*Dm*Cry), are blue-light photoreceptors through its FAD binding capacity.^12,13^ Type II Crys function as circadian clock regulators in a light-independent manner and do not bind FAD,^14–17^ rather they utilize the primary and secondary binding pockets as protein-protein interaction sites for the circadian proteins CLOCK and PER.^18–20^ Some studies suggest that FAD binding can be obtained through over-saturation of Type II Crys with FAD. ^21^ Type IV Crys, exemplified by avian Cry4a, bind FAD^16,22–24^ non-covalently and tightly. Substantial theoretical and experimental evidence supports that avian Cry4a undergoes photo-induced electron transfer between FAD and a conserved tetrad of tryptophans, a hallmark of radical-pair-based magnetic sensing *in vitro*.^8,16,23,25–28^

Despite the importance of FAD for the light-dependent functioning of the protein, the molecular mechanism of FAD binding in Cry proteins is only partially defined. In *Dm*Cry, single-point mutations hardly deplete FAD binding and double point mutations only partially deplete FAD binding by around 58%.^17^ Structural analysis suggests that FAD binding in *Dm*Cry is mediated through a cooperative effect of electrostatic interactions and steric modulation of the binding pocket accessibility.^29^ However, so far, there has been no killer experiment to completely deplete FAD from a Cry protein. Uncovering the FAD binding mechanism in avian Cry4 is therefore critical for elucidating the biophysical basis of magnetic sensitivity as it could possibly be manipulated directly through affecting FAD binding.

In this study, we use molecular dynamics (MD) simulations to identify key amino acids involved in FAD binding in the European robin (*Erithacus rubecula*) Cry4a (*Er* Cry4a). Fur-thermore, we generate a recombinant *Er* Cry4a mutant protein based on the computational results, and experimentally assess FAD binding ability using light absorption spectroscopy and mass spectrometry (MS). Finally, we determine the conformational difference between *Er* Cry4a wild type (WT) and mutant using native electrospray ionization mass spectrometry MS and ion mobility spectrometry MS, and propose a model of FAD binding mechanism in *Er* Cry4a.

We first analyzed interactions between *Er* Cry4a protein and FAD using Protein-Ligand Interaction Profiler (PLIP), a web tool for identification of non-covalent interactions between biological macromolecules and their ligands.^30^ The result suggests that 11 amino acid residues in *Er* Cry4a could interact with FAD (Figure 1A). The 11 amino acid redisues are S245, T246, T247, Q287, W350, H353, R356, F379, D385, D387, and N334. Strikingly, R356 is located parallel to both isoalloxazine and ribitol moieties of FAD, and interacts with FAD via diverse interaction including electrostatic-, cation-*π*, and hydrogen-bonding interactions. In contrast, S245, T246 and T247 appear to only form a weak hydrogen bonding network with FAD phosphate groups.

**Figure 1:**
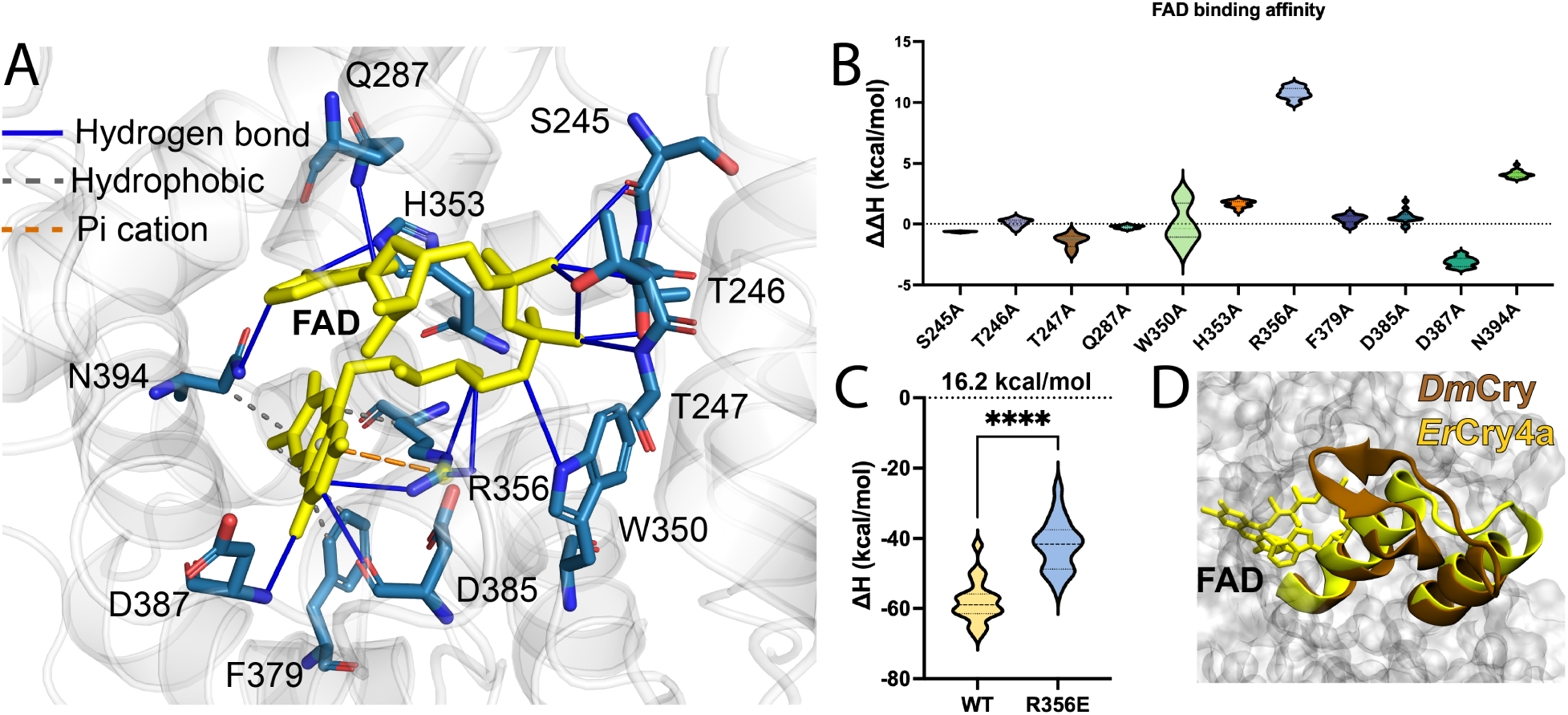
Computational simulation suggests key amino acids that affect FAD binding. (A) Illustration of interactions between amino acid residues and the flavin adenine dinucleotide (FAD) cofactor inside European robin (*Erithacus rubecula*) Cry4a protein (*Er* Cry4a). A total of 11 amino acid residues in *Er* Cry4a potentially interacts with FAD through hydrogen bond, hydrophobic interaction and cation-*π* interactions. (B) Predicted changes in the binding free energy upon mutation to alanine for the 11 amino acid residues that interact with the FAD. For the higher ΔΔH values, the mutations are predicted to more significantly disrupt FAD binding. (C) Predicted binding affinity for FAD in *Er* Cry4 wild type (WT) and R356E mutant. The absolute value of ΔH in R356E is 16.2 kcal/mol smaller than that of WT, indicating a weaker FAD binding affinity in R356E compared to that in WT. It should be stressed that the absolute ΔH values cannot be used to directly determine the FAD binding affinity, but these calculations reliably predict changes in binding affinities. (D) Structural comparison of *Er* Cry4a and *Drosophila melanogaster* cryptochrome (*Dm*Cry). The *β*-sheet on top of FAD in *Dm*Cry is absent in *Er* Cry4a, indicating a negligible steric constraints effect on FAD binding in *Er* Cry4a.

To quantify the interactions between the 11 residues and FAD, we computed binding free energy of X-to-alanine mutants, where X refers to one of the 11 residues. Alanine scanning mutagenesis is ideal for testing how much a particular residue contributes to binding because alanine is small and chemically neutral. ^31,32^ The computational alanine scanning result shows that the R356 residue stands out of the 11 amino acid residues surrounding FAD due to the high binding free energy (10 kcal/mol), indicating that R356 is the most energetically significant contributor to FAD binding (Figure 1B).

We hypothesize that the strong interaction between R356 and FAD is primarily driven by electrostatic forces, as arginine has a positive charge while FAD is negatively charged. To test the hypothesis, we introduced a charge-reversal mutation, replacing R356 with the negatively charged glutamic acid (R356E). Computational simulations suggest that the FAD binding affinity of R356E is 16.2 kcal/mol lower than that of the wild type, indicating substantially weaker FAD binding in the mutant (Figure 1C). Futhermore, structural analysis reveals that a *β*-sheet critical for FAD binding in *Dm*Cry is absent in *Er* Cry4a (Figure 1D), suggesting that steric constraints on FAD binding in *Er* Cry4a are negligible.

Following the *in silico* results, we generated the *Er* Cry4a R356E mutant in the wet-lab through site-directed mutagenesis and recombinant protein expression. During the gradient elution process of anion exchange chromatography, R356E mutant eluted from the anion exchange column at a solvent conductivity similar to that of WT protein (Figure 2A), suggesting that R356E mutagenesis maintains a similar surface charge pattern as in the WT protein. The purity of both protein samples is also similar, as shown by the single band on the Commassie blue staining image of a protein gel (Figure 2B). Most importantly, the UV–vis spectrum of *Er* Cry4a R356E mutant appears flat (near-to-null) in the absorbing range between 300 nm and 500 nm, indicating no blue-light absorption of R356E mutant and thus no FAD binding (Figure 2C). As a positive control, the WT protein shows vibrational fine structure around 350 nm and 450 nm, which are characteristic features typically associated with FAD binding in cryptochrome proteins.

**Figure 2:**
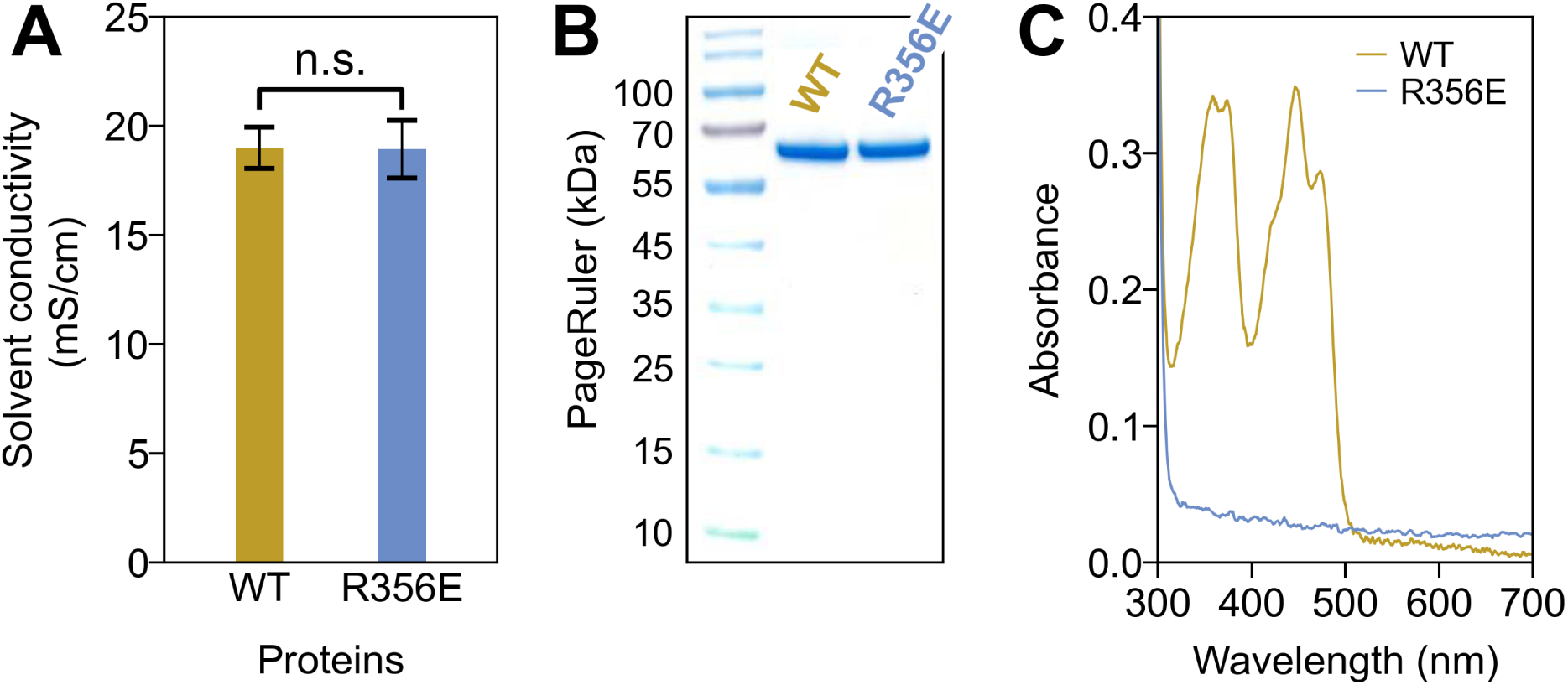
A single-point mutant *Er* Cry4a R356E completely deplete FAD from *Er* Cry4. (A) Both R356E and WT proteins elute from the anion exchange column at similar solvent conductivity during gradient elution process of anion exchange chromatography. (B) Sodium dodecyl sulfate – polyacrylamide gel electrophoresis Coomassie Blue Dye staining suggests that both proteins are comparably pure. (C) UV-visible absorption spectrum shows that the *Er* Cry4 R356E mutant does not absorb blue light between 350 nm and 500 nm, indicating the absense of FAD (the blue-light chromophore) binding. *Er* Cry4 wild type (WT) is used as a positive control showing characteristic spectral features of FAD binding.

Furthermore, mass spectrometry was employed to determine the protein sequence and cofactor binding status of *Er* Cry4a WT and R356E mutant. Under denaturing conditions, both proteins display broad charge state distributions, ranging from ∼ 18+ to 46+. The broad charge state distributions suggest that the proteins are completely unfolded and thus FAD was removed from the protein. This allowed assessing the subtle mass difference attributable solely to amino acid substitution. Deconvolution of the denatured spectra yields a mass difference of ∼ 29 Da between WT and R356E, which matches the theoretical mass difference between arginine and glutamic acid being 27.08 Da (Figure 3A). Consistently, a minor mass shift was also observed in native MS measurements on FAD-free WT protein and the R356E mutant protein (Figure 3B, the right panel). Together, the MS results validate that arginine has been successfully replaced with glutamic acid in the R356E mutant. Under native-like condition, *Er* Cry4a WT and mutant rotein charge state distributions narrows between 12+ and 16+, dominated by 13+ and 14+, consistent with folded conformations. The R356E mutant is detected exclusively without FAD, whereas the majority of WT (94%) binds FAD, indicated by a 786 Da mass difference (Figure 3A). It is interesting to note that, a dimer species is observed in both denaturing (Figure 3A) and native (Figure 3B) measurements. The protein dimers are very likely formed through the covalently disulfide bonds due to the absence of reducing reagent in the buffer solvent. The observation is consistent with our previous study on *Er* Cry4a dimer.^33^

**Figure 3:**
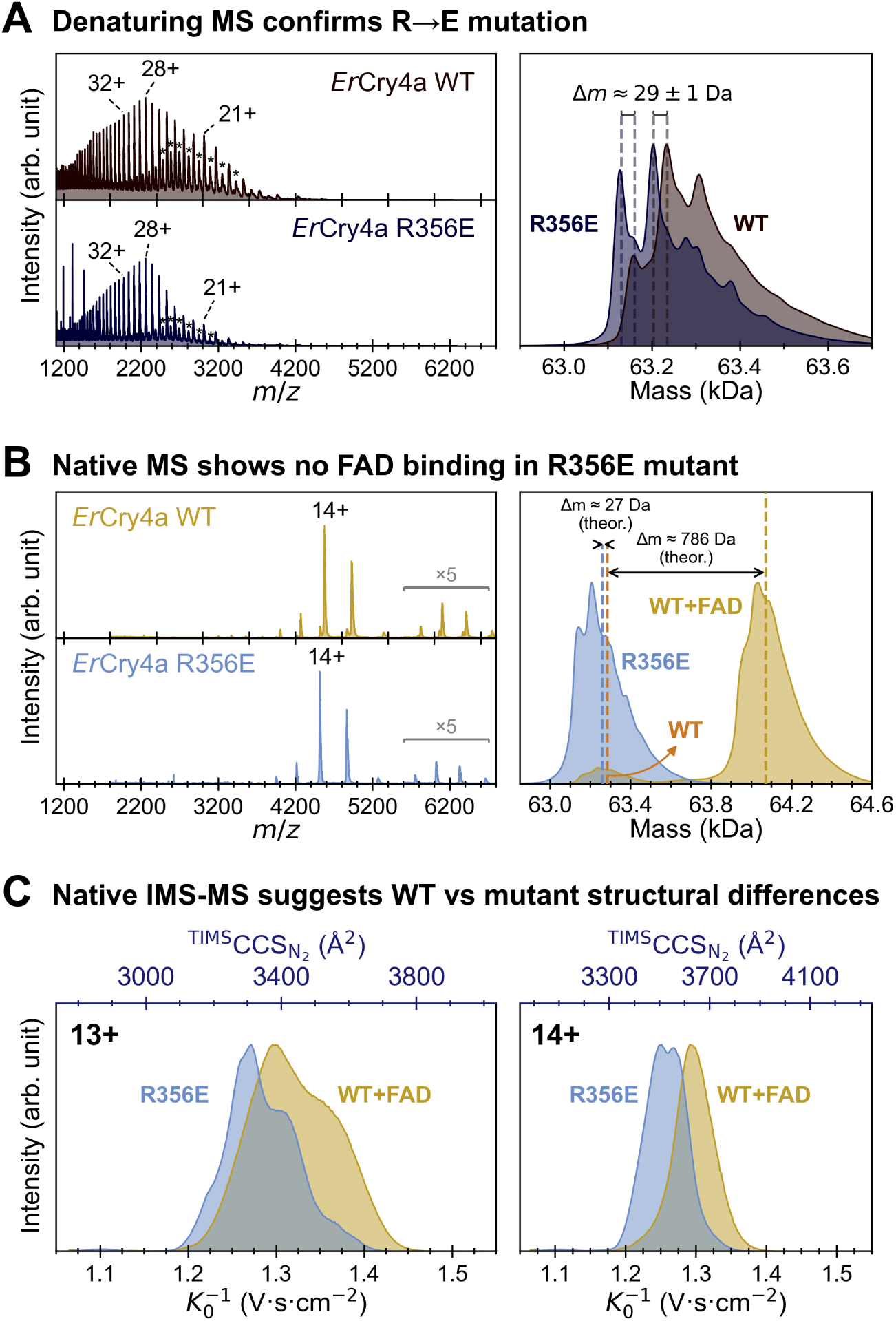
Mass spectrometry measurements validates that *Er* Cry4a R356E mutant does not bind FAD. (A) The denaturing MS detected a mass difference of ∼ 29 Da mass between WT and R356E mutant under the denaturing condition. The observed mass difference matches the theoretical mass difference between arginine and glutamic acid residues (27 Da), confirming the successful amino acid substition. The stars in the left panel mark denaturing protein dimers in the m/z range of 2400 and 3400. (B) Native MS detected a mass difference of 786 Da between WT and R356E under the native condition. The mass difference is consistent with the theoretical molecular weight of FAD. The dimer population is highlighted by a magnification factor of five at the m/z range of 5600 and 6800. (C) Ion mobility spectrometry - mass spectrometry (IMS-MS) shows that the collision cross section of R356E mutant is slightly left-shifted compared to FAD-bound WT protein, indicating a slighted more compacted conformation of R356E compared to the WT protein with FAD-bound.

Ion mobility spectrometry (IMS) measurements provide further insight into conformational properties of the two proteins. The FAD-bound WT protein exhibits a slightly larger collision cross section (CCS) value (Figure 3C) compared to the R356E mutant protein. For example, the CCS value at 14+ charge state is 3624.7 Å² for the FAD-bound WT protein and 3531.0 Å² for the R356E mutant protein. The measured CCS difference (ΔCCS) between the two protein is 2.7%. However, the expected CCS difference due to arginine substitution by glutamic acid is approximately ∼ 0.9%, which was calculated as CCS ∝ *m*^2/3^.^34^ The observed 2.7% shift therefore implies a conformational effect attributable to cofactor-dependent packing rather than the amino acid residue mass alone. Protein conformation of *Er* Cry4a R356E mutant is probably more compacted than that of the WT protein.

In summary, our study clearly shows that site-specific electrostatic repulsion is sufficient to completely block FAD binding in *Er* Cry4a. To our knowledge, this is the first report of a single-point mutation that completely depletes FAD binding in any Cry protein. Previous work on *Dm*Cry required dual mutations (R298E + Q311E) and the mutations only partially reduced FAD occupancy by 58%. ^17^ FAD binding in Cry proteins has been proposed to depend on two factors, i) non-covalent interactions between FAD and the binding pocket and ii) the steric availability of the binding pocket itself.^29^ In Type I Cry, strong non-covalent interactions and a conformationally closed binding pocket make FAD depletion difficult, typically requiring multiple mutations—one to weaken the non-covalent interactions and another to open the pocket.^17,29^ Specifically, in the *Dm*Cry double mutant, R298E was used to remove the *β*-sheet lid and Q311E was used to remove the pincer holding FAD in place.^29^ In the present study, we discovered that FAD binding is primarily mediated by electrostatic attraction, while steric constraints play a negligible role in *Er* Cry4a (Figure 4). It is perhaps suprising that one could achieve complete FAD depletion by a single-point mutation. However, an evolutionary divergence in the FAD-binding mechanism is likely to have occurred between Type VI Cry, exemplified by ErCry4a, and Type I Cry, exemplified by DmCry, due to differences in their protein structures. The apparent absence of a canonical β-sheet lid in *Er* Cry4a raises broader questions about how Cry proteins have structurally diversified to support species-specific physiological functions.

**Figure 4:**
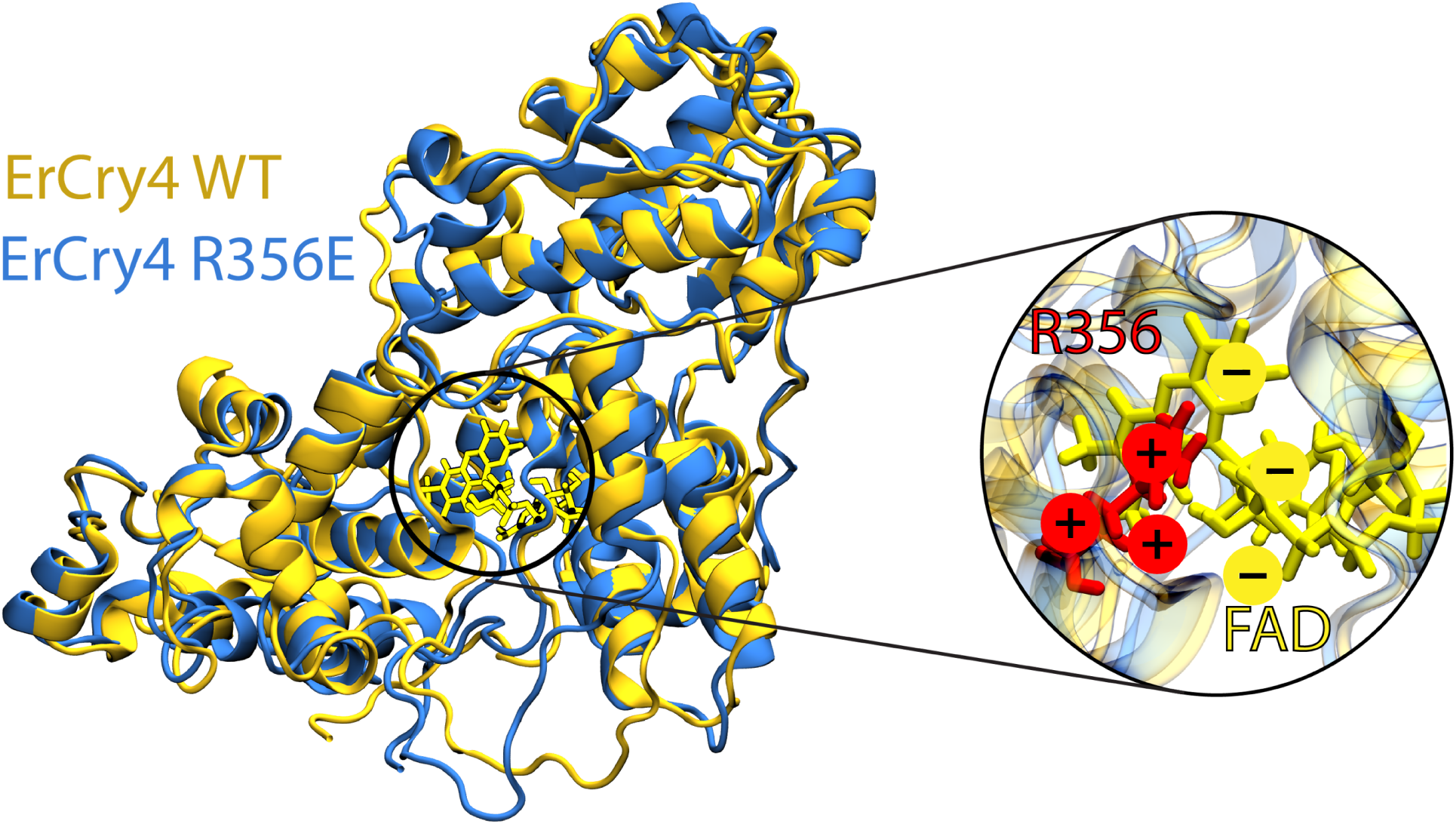
A model of FAD binding mechanism in *Er* Cry4a protein: electrostatic attractions secure FAD non-covalently binding in Cryptochrome 4a. The modelled structure of *Er* Cry4a WT and R356E mutant are shown in yellow and blue, respectively. The inset shows the interaction interface between the FAD and the R356E residue.

Single-point mutations are generally preferred in functional studies, as they introduce minimal perturbations to protein structure compared to dual or multiple mutations. In the present study, the single-point mutation preserved the overall charge profile of *Er* Cry4a WT protein, with only a slightly more compacted conformation observed in the mutant. The FAD-free R356E mutant of *Er* Cry4a will serve as an ideal negative control to investigate the structural and functional role of FAD in cryptochrome signaling and protein-protein interactions.^35^ In a much broader context, flavin-bound proteins have recently been employed as photosensitizers in photocatalytic proximity labeling, a technique that uses light-activated catalysts to tag nearby proteins or molecules for identification.^36^ A flavin-free Cry4a mutant can also provides a valuable negative control in this emerging application. In this regard, our study establishes a framework for generating flavin-free cryptochrome apoproteins.

In conclusion, this study provides new molecular insight into cofactor stabilization in avian Cry4a, revealing that electrostatic interactions, rather than steric constraints, dominate FAD retention. The R356E mutant serves as a powerful negative control for functional studies of *Er* Cry4a, enabling precise dissection of FAD-dependent signaling pathways in magnetoreception and in emerging photocatalytic proximity labeling applications. Future work should investigate whether the charge-reversal single-point mutagenesis strategy similarly disrupts FAD binding in Cry4a proteins from other bird species, and how FAD binding influences Cry4a conformational dynamics and protein–protein interactions, ^9–11^ which are both critical for understanding its role in light-dependent magnetic sensing.^7^

## METHODS

A homology model of *Er* Cry4a was built based on the *Cl* Cry4 template (PDB ID: 6PU0)^16^ in Swiss model workspace. The resulting *Er* Cry4a homology model was solvated in a water box with a padding distance of 12 Å. Hydrogens were added to the *Er* Cry4a model, while the topology was created using pdb2gmx from the gromacs software. ^37,38^ The protonation state of ErCry4a was estimated at pH 7 using PDB2PQR^39^ fromm the APBS biomolec-ular solvation software suite website. ^40^ FAD was parametrized using GAFF2,^41^ while the *Er* Cry4a protein was simulated using the ff99SB-ILDN forcefield.^3^ Specifically, simulations were carried out using a two femtoseconds timestep with the leap-frog integrator. Energy minimization prior to the molecular dynamics simulation was carried out using the steepest decent method. The temperature of the system was set to 300 K using the modified Berendsen thermostat and pressure was controlled at 1 bar using the Parinello-Rahman barostat.^42,43^ Long range electrostatic interactions were modeled using the Particle mesh Ewald summation approach.^44^ Constraints were set on hydrogen bond distances using the LINCS algorithm.^45^ Starting velocities were generated with a random seed. The simulation protocol included: i) energy minimization for 20,000 steps; ii) equilibration with restraints on the protein in the constant-temperature ensemble for 10 picoseconds; iii) equilibration in the constant-pressure ensemble for 100 picoseconds; iiii) production simulation in the constant-pressure ensemble for 50 nanoseconds. The residues that potentially influence FAD binding affinity were scanned using the protein ligand interaction profiler webserver.^30^ The *in silico* alanine scanning technique was subsequently used to quantify the contribution of the residues to the FAD binding affinity. Constant-pressure simulations were carried out following the equilibration and 20 frames were prepared as the input for the alanine scan analysis. Mutations of the FAD-surrounding amino acid residues to alanine were carried out using gmx_MMPBSA.^46^ FAD binding affinities for *Er* Cry4a wild type and mutant proteins were calculated using the Poisson–Boltzmann method implemented in gmx_MMPBSA.^46^ The analysis was carried out in Prism using the Welch’s *t* -test to find significance and differences in binding affinities.

*Er* Cry4a R356E mutant DNA was generated through a polymerase chain reaction (PCR) using the Q5 site-directed mutagenesis kit (New England Biolabs, Ipswich, MA, USA). The PCR primer sequences employed were 5’-TCACCTGGCTGAACACGCCGTCG-3’ and 5’-TGGATCCAGCCTTCCTGG-3’. The DNA sequence was confirmed via Sanger sequencing (LGC Genomics, Berlin, Germany). The *Er* Cry4a R356E mutant and the WT protein were expressed and purified as previously reported. ^23,25^ The UV-visible absorption spectra were measured using the Cary 60 UV-Vis Spectrophotometer (Agilent, Santa Clara, CA, US).

20 *µ*M protein was used for MS measurements. Prior to the native MS measurements, protein samples were buffer-exchanged to 150 mM ammonium acetate (pH 8) using ZebaTM Micro Spin desalting columns with a molecular weight exclusion limit of 40 kDa (ThermoFisher Scientific). Prior to the denaturing MS measurements, proteins were incubated in pH 2 150 mM ammonium acetate solution for 5 minutes. The MS and ion mobility spectom-etry–MS (IMS-MS) data were acquired on a timsTOF Ultra1 instrument (Bruker Daltonics) in positive ion mode using a CaptiveSpray 2 nanoESI source. To generate an ion source, the capillary voltage was set to 1.1 kV and dry gas was pumped at the speed of 3 L · min*^−^*^1^. The general temperature of the ion source was 80 °C. For the ion optics (native transmission), the parameters were as follows: Deflection 0 Delta = 105 V, Funnel 0 RF = 250 Vpp, Multipole 0 RF = 200 V, Deflection 1 Delta = 80 V, Funnel 1 RF = 400 Vpp, isCID Energy = 100-125 eV, Funnel 2 RF = 600 Vpp, Multipole RF = 1200 Vpp, Collision energy: 10 eV, Collision RF: 2000 Vpp, Ion Energy = 10 eV, Low Mass = 500 m/*z*, Tranfer time = 120 µs, Pre-pulse storage = 120 *µ*s.

The trapped ion mobility spectrometry in/out tunnel pressures used were 2.4/0.7 mbar (ΔP 1.7 mbar). The exact parameters were: Accumulation = 20 – 30 ms; 1*/K*_0_ ramp= 100 ms; 1*/K*_0_ window = 0.6 V · s · cm*^−^*^1^; scan ramp rate = 1.1 kV · s*^−^*^1^. DC potentials: DC potentials: Δ*t*_1_ = −20 V, Δ*t*_2_ = −120 V, Δ*t*_4_ = 100 V, Δ*t*_5_ = 0 V, Δ*t*_6_ = 20 − 40 V. m/*z* and 1*/K*_0_ were calibrated with ESI-L low-concentration tune mix (Agilent). The collision cross section (CCS) in N^2^ were calculated from the Mason–Schamp equation using the manufacturer CCS values. The MS and IMS-MS spectra were exported using Compass DataAnalysis software (Bruker Daltonics, v6.1). The deconvoluted spectra were obtains using UniDec^47^ with the following parameters: m/z ratio range: 1500-5000 Th, charge range: 10-50, mass range: 30-150 kDa, sample mass set to every 5 Da, peak FWHM: 2 Th, peak shape: Gaussian.

## Author Contributions

J.X. and E.S. contributed equally to this work.

## Notes

The authors declare no conflicts of interest.

## Acknowledgement

This work was supported by Lundbeck Foundation (Postdoc fellowship grant number: R402-2022-1372, awarded to J.X. grant number: R303-2018-3175, awarded to E.S.R); Novo Nordisk Foundation (INTEGRA, grant number NNF20OC0061575 to O.N.J.); European Research Council (under the European Union’s Horizon 2020 research and innovation programme, grant agreement no. 810002, Synergy Grant: ‘QuantumBirds’, awarded to H.M.); Volkswagen Foundation (Lichtenberg Professorship to I.A.S.), the Deutsche Forschungsgemeinschaft (SFB 1372: Magnetoreception and Navigation in Vertebrates, no. 395940726, and EXC-3051: NaviSense, no. 533653176 to H.M. and I.A.S); TRR386/1-2023 Hyperpolarization in molecular systems HYP*MOL, no 514664767 to I.A.S.), the Ministry for Science and Culture of Lower Saxony (Simulations Meet Experiments on the Nanoscale: Opening up the Quantum World to Artificial Intelligence (SMART) and Dynamik auf der Nanoskala: Vonkohärenten Elementarprozessen zur Funktionalität (DyNano)).

---

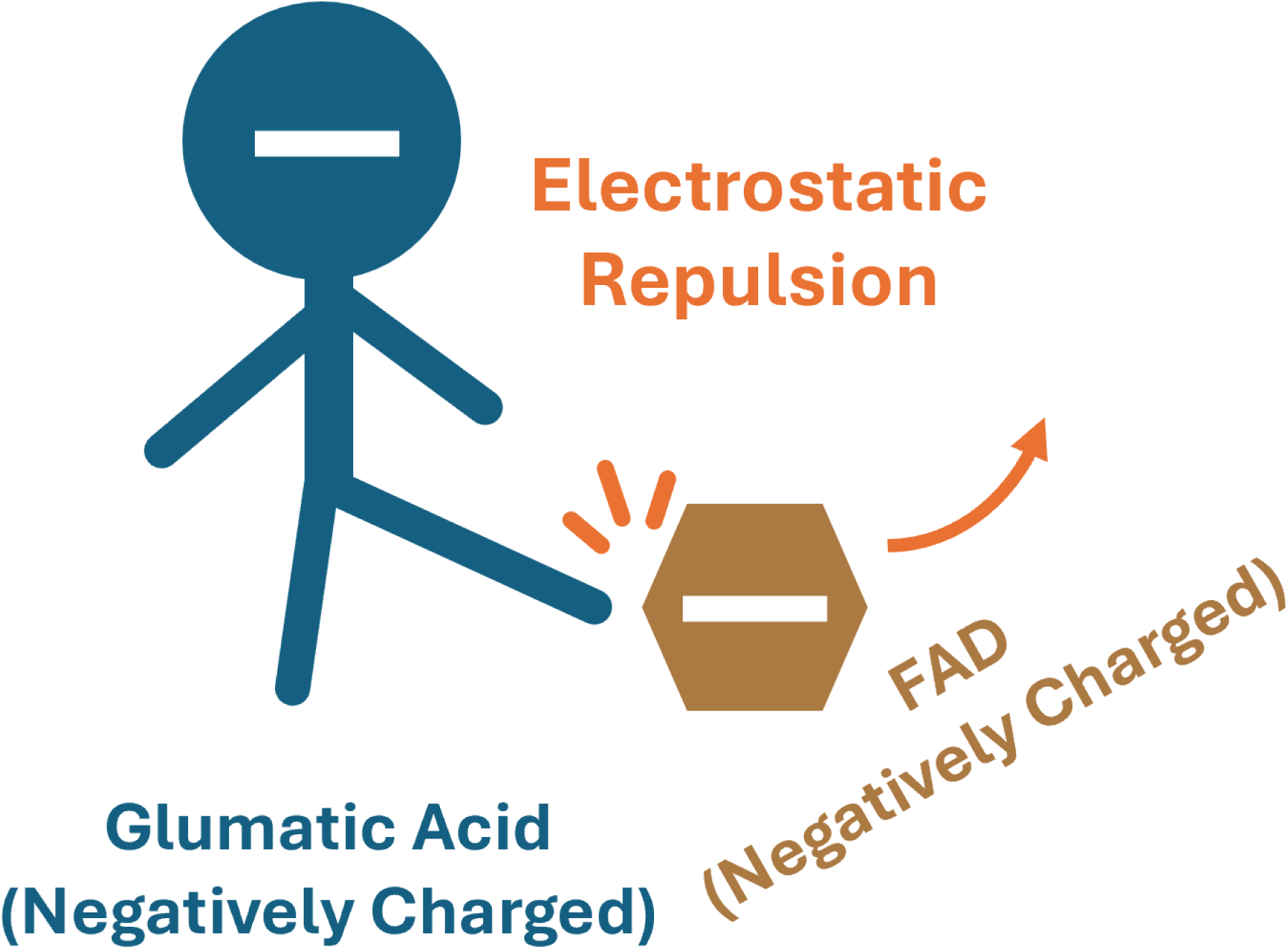

